# REM-like Oscillatory Theta Activity Predicts a Reduction in the Recuperative Value of Natural Human Sleep

**DOI:** 10.1101/2023.10.06.561209

**Authors:** Shashaank Vattikuti, Tracy J Doty, Samantha Riedy, Allen Braun, Thomas J Balkin, John D Hughes

## Abstract

Here, using data from two independent studies, we examine whether all of sleep is restorative or paradoxically whether some sleep processes incur a sleep debt that impacts next-day wakefulness. Specifically, we examine whether rapid eye movement (REM) sleep is such a process due to its similarity to wake activity, which is causal for sleep debt. To investigate this, we first develop a novel measure of REM neural activity (REM-like oscillatory theta activity (OTA)), overcoming limitations of current sleep scoring. We find that naturally occurring average REM-like OTA across individuals: 1) is associated with increased neurobehavioral sleep debt; 2) explains 25-38% (p ≤ 0.001) of sleep debt differences across individuals the following day; 3) occurs throughout sleep to various degrees, contrary to current sleep scoring; and 4) can be measured automatically, without cumbersome manual scoring.

## Introduction

Human sleep is dynamic, with multiple, substantial changes in EEG activity over the course of a normal nighttime sleep period. Sleep is generally considered to provide recuperative brain processes that enhance subsequent waking alertness. The EEG is an obvious modality to monitor this recuperation. However, underlying sleep signals in the EEG are the product of or support a complex combination of sleep processes (e.g., elimination of metabolic waste products/neurotoxins from the parenchyma; replenishment of ATP and energy sources such as glycogen; reduction of adenosine, cytokines, and other sleep-regulatory substances; synaptic plasticity^1–5^), any of which may or may not contribute directly to the overall recuperative function of sleep. In fact, it is conceivable that some sleep processes may even incur additional “sleep debt”. Sleep debt is a hypothetical construct that lacks a well-accepted, singular physiological definition but its existence is inferred from a multitude of measures (e.g. subjective fatigue, multiple sleep latency tests, maintenance of wakefulness tests, cognitive tasks, cardiovascular measures). More narrowly, sleep debt is considered to be primarily dissipated by sleep processes with associated slow-wave activity (SWA) that predominates the first several hours of nighttime sleep. Rapid eye movement (REM) sleep is a likely candidate for a constellation of sleep processes that, in sharp contrast to SWA, actually incur additional sleep debt – a possibility that is the main focus of the present study. Specifically, we hypothesized the following: 1) that the *REM-like* neurophysiological components of REM sleep (that may or may not strictly co-occur with other conventionally-defined REM sleep phenomena such as eye-movements and atonia) incur a sleep debt similar to wakefulness, 2) REM-associated oscillatory theta-band activity (OTA) (described later) reflects this sleep debt-incurring process, and 3) these effects are relevant to natural human sleep and impact subsequent wakefulness. [We define “natural” as conditions with sleep dynamics that are not altered by direct interference (sleep interventions such as experimentally induced awakenings, medications, etc.)] Prior studies have shown that REM sleep is important for nervous system development^6^, neural plasticity^7^, and the processing of emotional memories^8^. We investigated whether the beneficial functions of REM sleep physiology come at a cost to recuperation, as reflected in next-day measures of alertness.

This *REM-OTA Fatigue Hypothesis* is premised on the presence of those wake-like EEG features that characterize REM sleep and other shared neurophysiology. These wake-like features include activity levels in those EEG frequency bands (theta, beta, gamma) that are thought to support conscious cognitive functions^9–11^, and elevated cerebral blood flow levels that are suggestive of greater energy demand^12,13^. It has previously been shown that activity during REM sleep in these bands reaches levels similar to (or even greater than) those evident during wakefulness, and that they occur in conjunction with higher levels of acetylcholine (necessary for conscious information processing^14,15^) than during either non-REM (NREM) sleep or wakefulness^16,17^. Although *REM-OTA Fatigue* provides a compelling hypothesis, there is a dearth of studies that address this hypothesis during natural human sleep. A major obstacle to testing this hypothesis has been uncertainty about how to measure the intensity of REM-like neurophysiology under natural conditions. This specific issue has been expressed by Peter Achermann, a major proponent of classic sleep models (2002 & 2023, personal communications).

Accordingly, we addressed this question (and thus the second, OTA, component of our hypothesis) by investigating the extent to which *REM-associated oscillatory theta-band* (4-8Hz) activity (*REM-like OTA*) is predictive of subsequent daytime neurobehavioral sleep debt. We adopted this approach to circumvent well-known limitations of conventional sleep staging^18^ and raw spectral band power. Notably conventional clinical sleep staging (scoring) using the American Academy of Sleep Medicine (AASM) guidelines, the global clinical and research standard, grossly under-utilizes the richness of EEG recordings. Importantly, it has been suggested that NREM and REM sleep processes are not as temporally dissociable as traditional sleep scoring implies. REM physiology may be present to a degree during sleep periods otherwise scored as NREM (referred to as “covert REM”)^19^ and conversely, NREM features such as slow-wave activity may occur during REM periods^20,21^. In fact, various measures of “REM-like” neural activity outside of conventionally-scored REM periods have been recently explored using invasive methods in non-human animal models^22^. As we describe below, raw spectral power of the sleep EEG is inadequate for measuring specific neurophysiological processes even with the aid of manual scoring (in part due to confounding from multiple sleep processes) and more targeted signal processing is warranted.

To non-invasively quantify REM-like sleep neurophysiology in humans in a manner agnostic to conventional sleep stage scoring, we used measures of REM-like OTA with scalp-EEG that we previously developed^23^ (see *Methods*). This decision was based on the presumed physiological relationship between superficial EEG *REM-like OTA*^24^ and *hippocampal generated theta activity* (which is an essential component of hippocampal memory processes during wakefulness in mammals and is frequently present at high amplitude during REM sleep in most species studies; suggestive of a causal role in memory functions during sleep and supported by at least some intervention studies^25,26^) or cortical activity with a similar function ^17,27,28^. How to specifically measure REM-like oscillatory theta activity using non-invasive scalp EEG is an open question since, as we show, raw spectral theta-band power is composed of multiple processes. The modifying term “oscillatory” in “OTA ’’ signifies a difference from arrhythmic (1/f) background activity^29^ that is believed to be functionally unrelated to this hippocampal-related theta and is much higher in power. We hypothesized that oscillatory theta activity would be an accurate reflection of those aspects of REM physiology that incur sleep debt because: 1) theta power is associated with significantly increased power in higher frequency bands (beta, gamma) in REM sleep that are metabolically demanding^13^ and 2) theta activity is directly implicated in waking and REM sleep related plasticity processes^17,30^. Both phenomena are thought to be major components of the process by which homeostatic sleep drive accumulates^31^. The term “REM-like” is meant to reflect the fact that this activity is characteristic of conventionally-scored REM sleep, but may also be present (albeit to a lesser extent) during conventionally-scored NREM sleep^19,32^. To address this we leveraged a fractal theory (the IRASA process^33^) to isolate REM-like oscillatory theta from 1/f aperiodic activity. Then we introduce a normalization to isolate REM-like OTA from other sources of oscillatory theta and nonparametric filtering to reduce sample noise. In this way, we derive a time-activity series for REM-like neural activity across the nighttime sleep period.

As a test of the *REM-OTA Fatigue Hypothesis*, we assessed the association between prior night REM-like OTA and next-day neurobehavioral sleep debt using the current standard for this, the psychomotor vigilance test (PVT). As noted above, while the physiological definition of sleep debt is debatable, numerous cognitive and subjective measures are impacted by sleep loss to varying degrees. A challenge for sleep studies that involve severe sleep loss (sleep deprivation or chronic sleep restriction) is administering a sensitive, rapid, and simple assessment for cognitive impairment. The PVT has stood out as such a test and is considered the gold-standard measure for neurobehavioral sleep debt. It is a reaction time test, where an individual simply responds to the appearance of a randomly presented cue. Sustained attention (vigilance) as assessed by the PVT is exceedingly sensitive to sleep loss^34^ as well as the circadian rhythm of alertness^35,36^. It is well correlated with other cognitive tests such as Digit Symbol Substitution^37,38^ and the Serial Subtraction/Addition Task^38^, and to a somewhat lesser extent with measures of subjective sleepiness. The degree to which it deviates from subjective measures is largely thought to reflect a relative lack of sensitivity of those subjective measures^37,39–41^. It is used in many contexts as a research tool for assessment of sleep loss-induced cognitive impairment including by the military^42^ and NASA^43^, and in the assessment and quantification of driver fatigue^44,45^. In addition, the extrema of PVT performance have been found to be predictive of mental resilience/vulnerability to sleep loss across individuals^46,47^. Finally, PVT attentional lapses and slowed response times (impairment) have been shown in several studies to increase along with theta- and delta- (0.5-4Hz) band power during extended wakefulness^46,48^. In this case, unlike the REM-like OTA measure used here, this band power is thought to reflect homeostatic sleep drive^49^ (sleep debt) during wakefulness. The PVT, in addition to being well validated, is currently a more tractable assessment tool for incurred sleep debt than EEG and was employed in each of the sleep studies used in the present analyses.

In short, we evaluated the REM-OTA Fatigue Hypothesis by testing whether the average whole night REM-like OTA correlates negatively with PVT performance (thus suggesting that REM-like physiology does indeed result in the accrual of sleep debt). We tested this hypothesis using data from two independent sleep studies (whose primary objectives have already been presented^42,50^) in which sleep schedules were designed to minimize sleep debt acquired during the wake period across all individuals. This was accomplished by utilizing sleep EEG data collected under conditions of normal to extended sleep durations (aka “sleep satiation”) that preserved natural sleep processes. Logically, this sleep satiation minimized sleep debt effects and thus increased the likelihood that alertness deficits incurred by REM OTA would be detectable.

## Results

### REM-like oscillatory theta activity (OTA) during scored-REM is quantitatively associated with reduced next-day vigilance

First, we determined the extent to which REM-like OTA during conventionally scored REM sleep (i.e., using AASM guidelines, hereafter referred to as *scored- REM*) epochs was predictive of next-day PVT impaired-performance (inter-individual differences in mean reaction times, *PVT-RT*). As shown in Figure 1a, there was a substantial and statistically significant positive correlation (n = 42, r = 0.5, p = 0.001) between REM-like OTA and PVT-RT across individuals using the combined data from the two studies. Higher REM-like OTA was associated with poorer subsequent next-day performance (longer RTs). Prior night REM-like OTA explained 25-38% of the inter-individual variance in next-day impaired- performance. The upper bound was estimated by removing one major outlier shown in Figure 1a (n=41, r=0.63, p<0.001). Figures 1b-c show that this association was not an artifact produced by combining the data since the effect was also evident when data from each study was analyzed separately. (All subsequent analyses used the combined data from both studies.)

**Figure 1.**
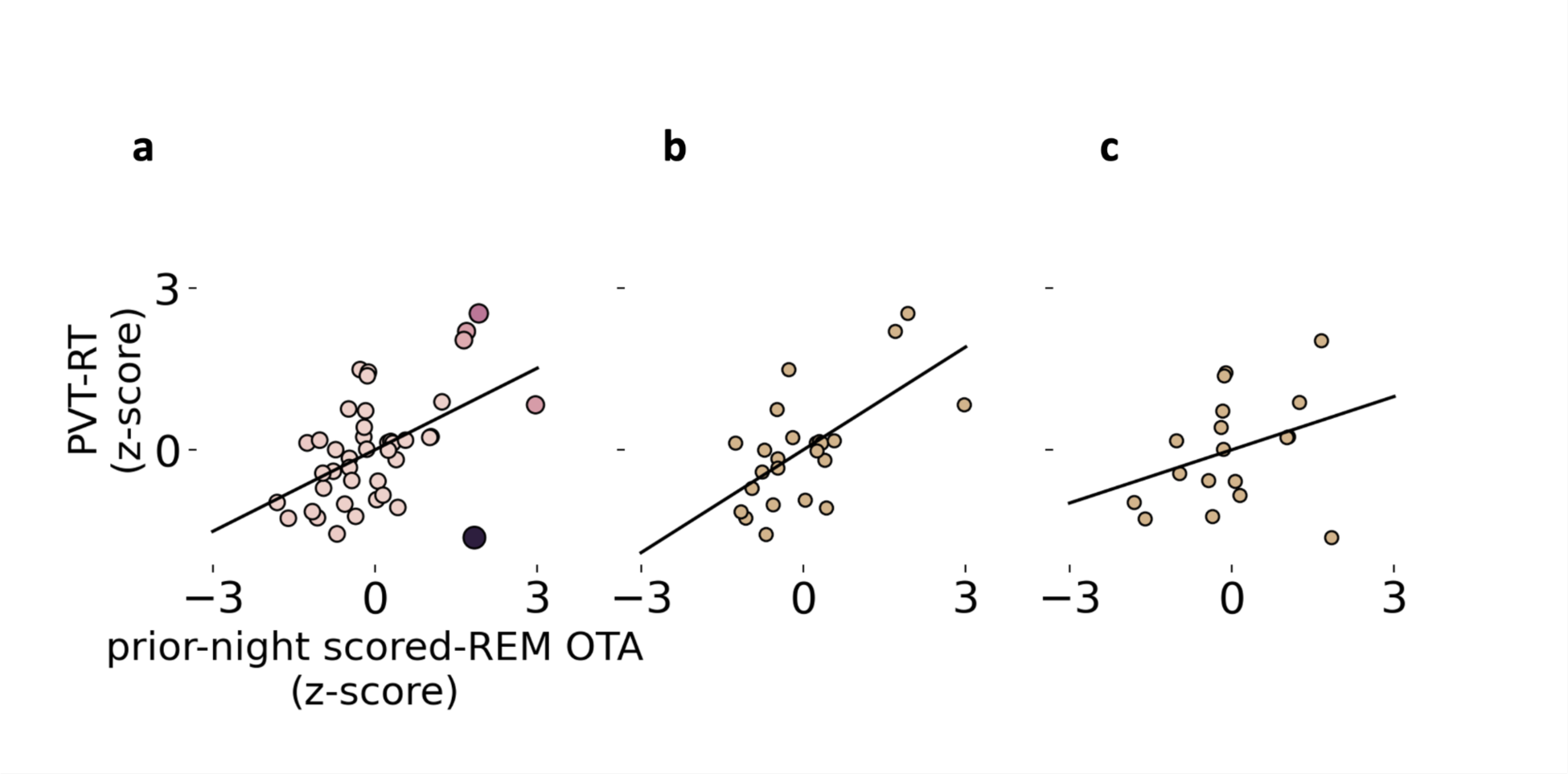
Correlation between prior whole-night scored-REM oscillatory theta and next–day impaired-performance (PVT-RT). Results from two independent studies. (**a**) Combined study correlation: n = 42, r = 0.5, p = 0.001 (removing one outlier based on Cook’s distance > 0.5, shown in black: n=41, r=0.63, p<0.001). (**b**) Study A results : n = 24, r = 0.64, p = 0.001, (**c**) Study B results : n = 18, r = 0.33, p = 0.17. PVT-RT is the next-day average response time for an individual across all sessions, which were z-scored across individuals in both studies. Prior-night scored-REM OTA is the IRASA-derived total 4-8Hz band “oscillatory” power across all manually scored REM epochs (whole-night spectra). These were also z-scored for each study. The regression line for each condition is shown in black. For the combined study regression (**a**), outliers were estimated using Cook’s distance indicating leverage of each data point, shown by hue and size.

### Under sleep satiated conditions, standard sleep measures fail to predict next-day vigilance but REM measures consistently predict reduced performance

Next, we compared the strength of the association between REM-like OTA during scored-REM and next-day performance impairment (PVT-RT) versus the association between more traditional (standard) sleep parameters and next-day PVT-RT. Standard measures included total sleep time (TST) (which varied only as a function of sleep efficiency - proportion of sleep vs wake during the sleep period - since time in bed was held constant in each study), relative scored-N3 duration (to all sleep epochs) (“deep sleep”), relative scored-N2 duration (“light sleep”), relative scored-REM duration, average slow-wave activity (0.5-4Hz) power, and average raw scored-REM theta-band power (again “scored” refers to conventional AASM scoring methods). None of these standard sleep measures were significantly associated with next-day PVT-RT (see Figure 2). Lack of a TST effect suggests that sleep satiation was achieved by the study designs; i.e., individual sleep needs due to prior wakefulness were largely satisfied. The lack of SWA effect may be due to a similar reason but also SWA, while often investigated, has not been found to be a robust predictor of next-day sleep debt.

**Figure 2.**
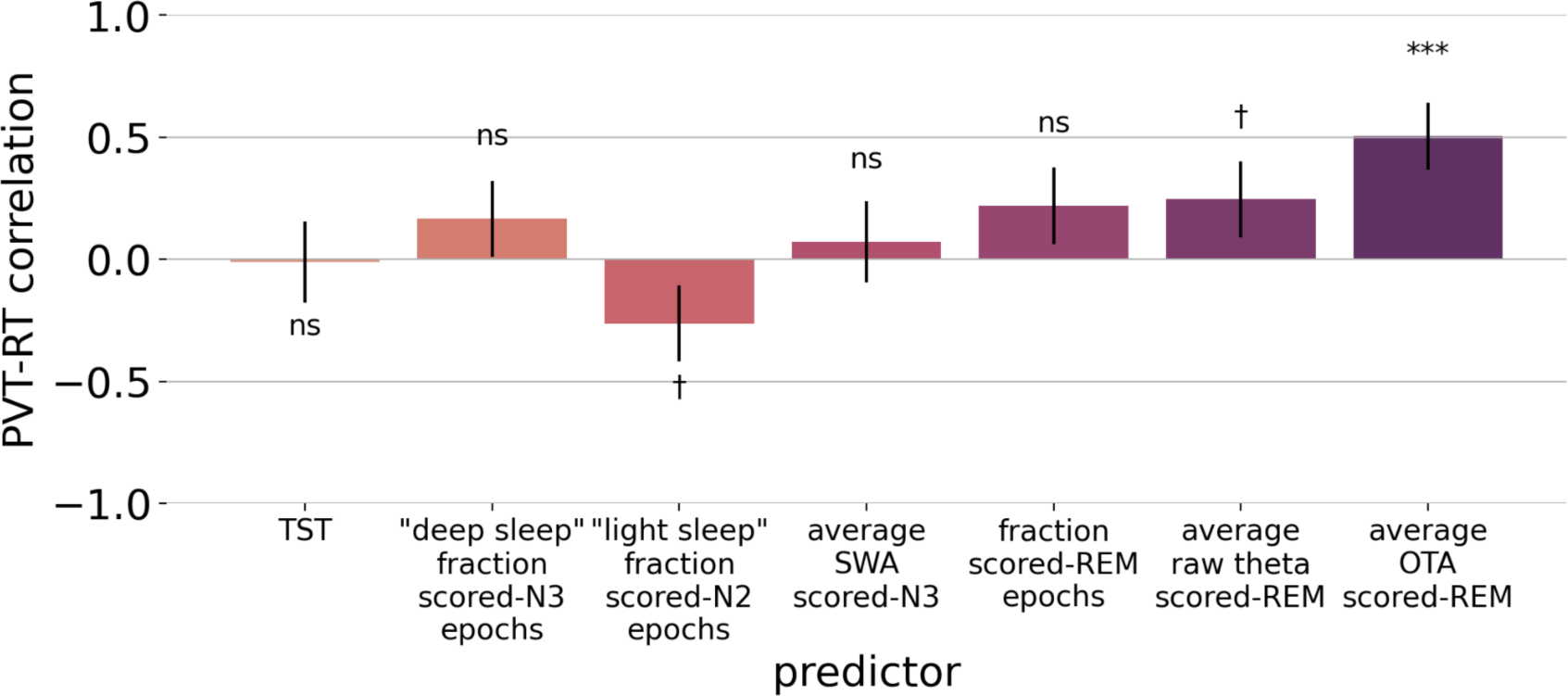
Scored-REM OTA vs conventional sleep measures as predictors of next-day impaired-performance (PVT-RT). Key: TST=total sleep time, “deep sleep” = ratio of N3 epochs to total sleep epochs, “light sleep” = ratio of N2 epochs to total sleep epochs, REM epochs = ratio of REM epochs to total sleep epochs, SWA = average raw 0.5-4 Hz power across N3 epochs, raw theta = average 4-8 Hz power during REM epochs, and OTA=average oscillatory theta activity 4-8Hz during REM epochs (***p<0.001, **p<0.01, *p<0.05, † p≤0.1, ns p>0.1).

In sharp contrast, all REM measures showed a consistent positive correlation with next-day PVT-RT. Also, the effect size (correlation) increased as the measures became more specific for OTA, supporting that REM neural activity was the underlying factor (relative conventionally scored-REM sleep duration< average raw theta-band power during scored-REM<average OTA during scored-REM).

### Covert REM (REM-like OTA during scored-NREM) is quantitatively associated with reduced next-day vigilance, whole-night vs per-epoch

Next, we determined the extent to which covert REM is also predictive of next-day impaired- performance. For this analysis we replicated the previously described analyses using only AASM scored- NREM epochs. Although the sign was consistent (positive association between scored-NREM REM-like OTA and performance impairment), in our initial analysis the signal was noisy and failed to reach statistical significance (N2 r=0.2, p=0.2; N3 r=0.21, p=0.17). We hypothesized that this was due to multiple sources of oscillatory theta-band activity during NREM periods - i.e., sources that are not REM-like OTA. For example, transitions to the hyperpolarized phase of the slow oscillation (SO) (the traditional, electrophysiological hallmark of slow wave sleep and thought to be maximally recuperative) can be preceded by theta-band bursts^51^. Also, REM-like OTA, if it exists during scored-NREM 30-s epochs, is likely to be less frequent and weaker, and thus harder to detect.

In a prior analysis^23^, we found that adding a per- (30 second) epoch normalization factor (see *Methods*) helps to isolate REM-like OTA signals without using conventional scoring. All analyses above used “whole-night” data, in which all manually scored epochs corresponding to the target stage were concatenated and spectral measures were acquired across a much longer time series than a single epoch. However, to run the normalization we needed to perform this on each epoch to produce a weighted average across scored-NREM epochs with more or less REM-like OTA. This per-epoch IRASA introduces more finite sample noise, so we also used a nonparametric filter to reduce spurious signals. Scored-NREM epochs were selected prior to performing the filtering to reduce contamination from scored-REM epochs.

As shown in Figure 3, the normalized per-epoch REM-like OTA (nOTA) strengthened the PVT-RT associated signal during scored-NREM periods (N2 r=0.37, p=0.016; N3 r=0.23, p=0.14) with a pronounced effect in the N2 covert REM signal. There was also a small decrement for scored-REM periods (r=0.38, p=0.01). (Note that there is a negligible discrepancy between scored-REM REM-like OTA estimates in Figure 1 and 3 due to differences in the averaging method used to maintain consistency with the normalized per-epoch OTA analysis.) We also assessed the extent to which normalization was driving these associations, but found no significant association between the normalization and next-day impairment for any of the stages (p-values between 0.2-to-0.8). This is depicted in Figure 1 which shows that during scored-REM, REM-like OTA without normalization was found to be a good predictor of next-day performance (presumably normalization was not needed due to a stronger REM-like OTA signal during scored-REM compared to scored-NREM epochs).

**Figure 3.**
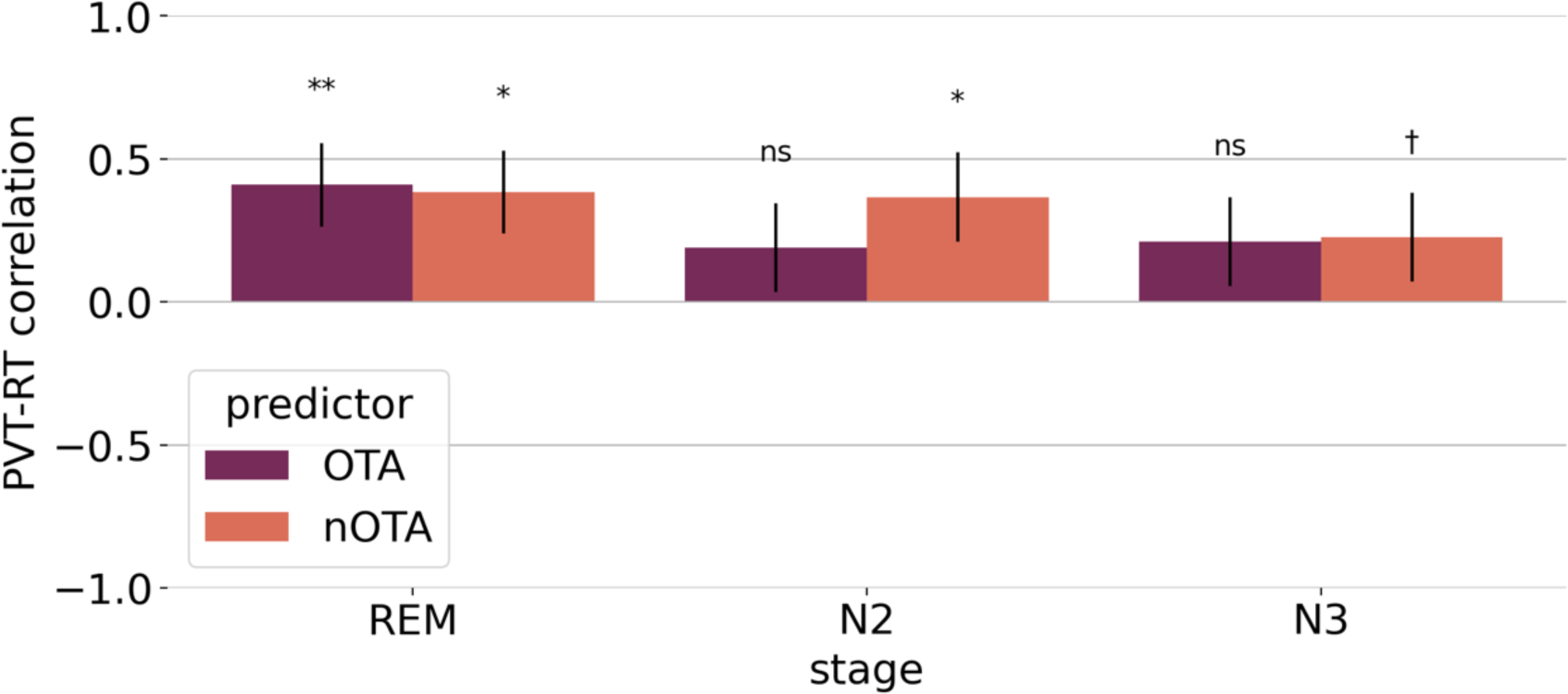
Covert REM next-day impaired-performance effects. Shown are the correlations between the average oscillatory theta activity (OTA) and normalized OTA (nOTA) for different conventional stages and next-day mean PVT reaction time (PVT- RT) (***p<0.001, **p<0.01, *p<0.05, † p≤0.1). There is a slight discrepancy between Figure 3 scored-REM OTA and Figure 1, due to different averaging using the per-epoch IRASA derived measures.

### nOTA is an inherently automated, conventional-score agnostic measure of REM-like neural activity

The aforementioned analyses utilized sleep scoring data based on AASM criteria. However, since the REM-like OTA and PVT association was also evident during scored-NREM epochs, we hypothesized that the observed relationship between REM-like OTA and next day performance may be detectable without standard sleep scoring. This was considered desirable given the well-known issues with standard manual sleep scoring. As shown in Figure 4 and noted above, both REM-like OTA and nOTA during scored-REM were significantly correlated with impaired next-day performance (scored-REM, REM-like OTA r=0.4, p=0.006; nOTA r=0.38, p=0.01). Most importantly, this correlation was also evident when sleep stage scoring was discarded (labeled “All” in Figure 4) [All-epochs including wake-after-sleep-onset (WASO) REM-like OTA r=0.3, p=0.047; nOTA r=0.52, (p<0.001) - these analyses utilized all epochs during the time-in-bed (TIB)]. The correlation with PVT-RT was strongest with nOTA, although both measures were significantly correlated with performance. Because it was possible that WASO, due to sleep debt accruing wake periods, contaminated the TIB analysis, we repeated the analysis with all scored wake epochs removed. The correlation was still significant (n=42, r=0.5, p<0.001), indicating that the sleep stage-agnostic results were unlikely to be an artifact of wakefulness during the sleep period.

**Figure 4.**
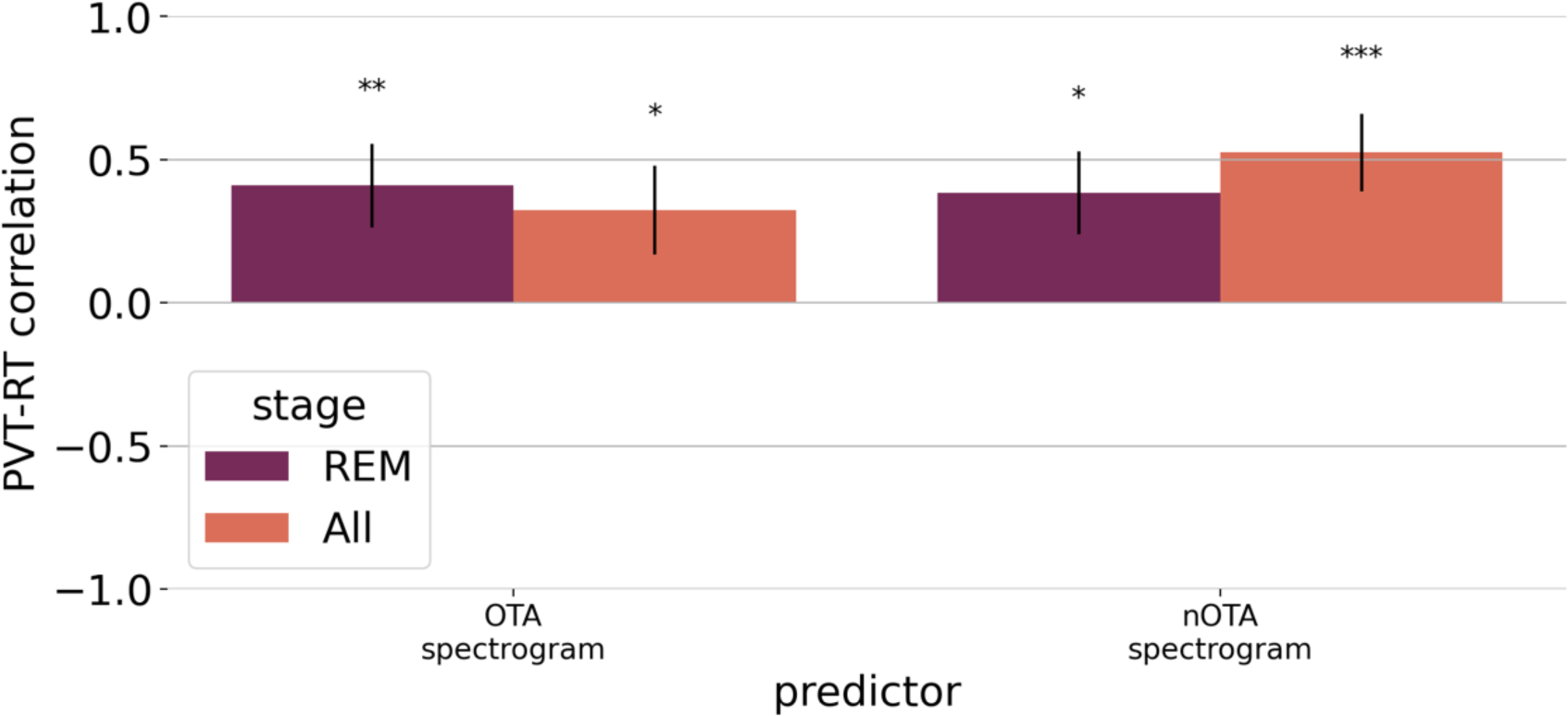
AASM scored-REM vs AASM-agnostic (All including wake-after-sleep-onset) oscillatory theta activity (OTA) and normalized OTA (nOTA) as predictors of next-day impaired-performance. (***p<0.001, **p<0.01, *p<0.05, † p≤0.1)

As illustrated in Figure 5 (adapted from our prior study^23^), nOTA is a superior quantitative measure (compared to AASM sleep scoring methods) for tracking REM-like activity. Raw theta-band activity mostly reflects the NREM stage 3, 1/f-type signal. As shown, REM-like OTA is more specific to scored- REM epochs. However, the nOTA measure appears to be even more specific to REM-like activity. In fact, the presence of this REM-like activity during traditionally-scored NREM epochs can be considered a form of “covert REM”, or mixed REM/NREM physiology, as evidenced by the preceding analyses. nOTA is a quantitative, automated, physiologically interpretable, and outcome-validated (as reflected in the present work) measure that accounts for REM neural activity across both scored-REM and NREM epochs.

**Figure 5.**
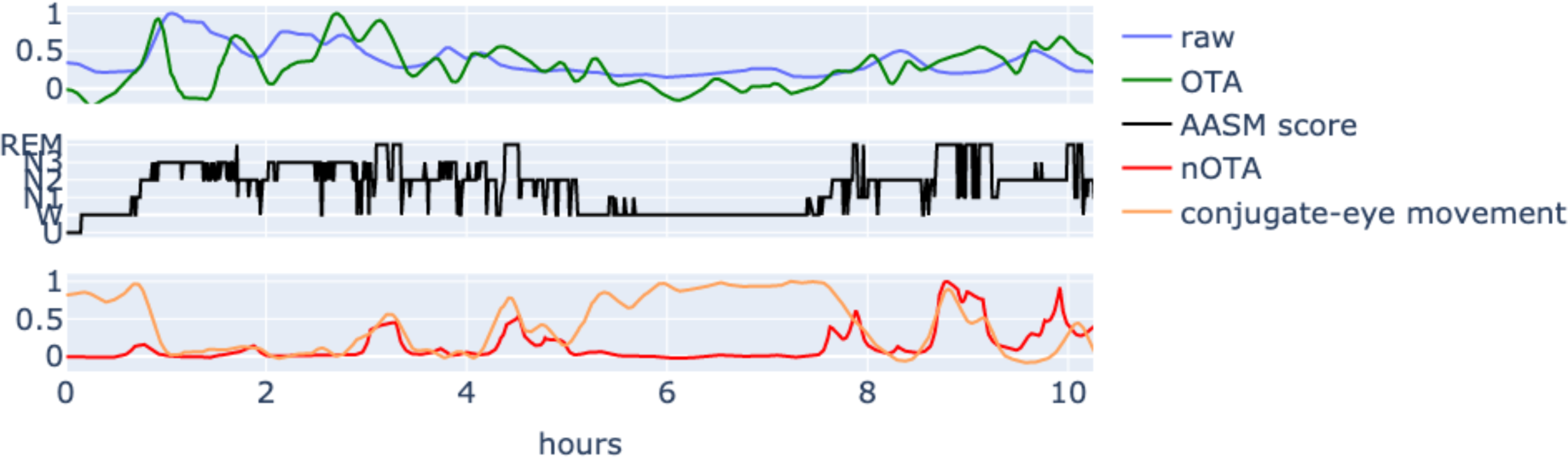
Theta-band measures across the sleep period for a single individual. A visual depiction of the extent to which different measures of 4-8Hz band activity track sleep stages and conjugate eye movements (associated with REM sleep and wakefulness) across the night (see *Methods*). All sleep measure traces in this figure are scaled to their maximum amplitude to facilitate visual comparisons.

### Oscillatory nature of the REM-like activity

Visual inspection of the IRASA spectra (Figure 6) is consistent with oscillatory activity in the theta-band range (4-8 Hz). (These results are intended to be descriptive of the oscillatory nature with no formal hypothesis testing.) Taking the median spectra across subjects, the IRASA-inferred oscillatory power was clearly elevated in the standard 4-8 Hz band for both studies. We observed a clear peak that spanned the 4-10 Hz range. A closer look at individuals, exemplified in Figure 6b, suggests that the extension of the group-wise peak into the standard 8-13 Hz alpha range is due, at least in part, to the presence of both oscillatory theta- and alpha-band activity with varying levels of visually distinct (separate) peaks. This is supported by the observed differential power between frontal (Fz) vs posterior (O1) channels and REM vs WAKE conditions. For example, Subject-0 shows two peaks. The lower frequency peak, we presume is oscillatory theta (OTA) and the higher, oscillatory alpha. This is supported by the higher power during REM of the OTA peak in the frontal channel compared to the posterior, occipital channels and dominance of the presumed alpha peak during wakefulness. In addition, some individuals such as Subject-1 have OTA peak frequencies around 8 Hz. These figures support our use of the 4-8 Hz bandpass in the above regression analyses to capture OTA and reduce oscillatory alpha signal. While further optimization may be warranted, the main regression results clearly support the utility of isolating such oscillatory-like dynamics.

**Figure 6.**
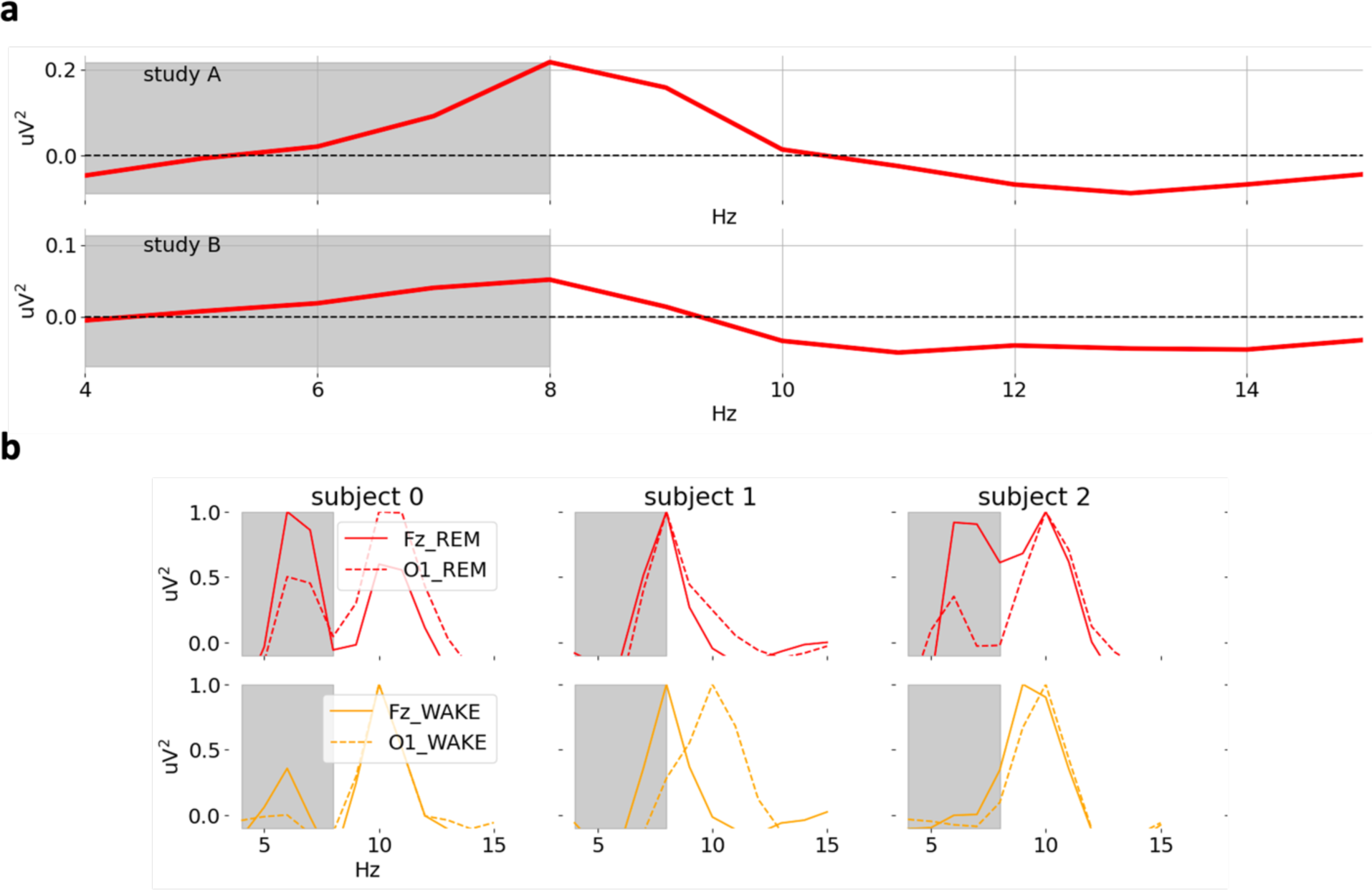
IRASA-derived oscillatory spectra for AASM-scored REM. Inferred oscillatory spectra using the IRASA method. (**a**) Average (median) power spectra across individuals for AASM-scored REM (red) epochs, channel Fz (mastoid reference) (solid). (**b**) Example spectra of individuals. Contrastive plots between frontal (Fz) and posterior (O1) (dashed) channels and scored-REM vs scored-WAKE (orange). Gray-shading indicates the conventional theta-band used for the regression analyses (4- 8 Hz). Figure intended for descriptive purposes only with no formal hypothesis testing.

## Discussion

The present findings are consistent with the hypotheses that: 1) REM sleep physiology during natural human sleep incurs a sleep debt that impacts subsequent wakefulness, and 2) oscillatory theta activity - a feature of REM-like neurophysiological activity that occurs during both AASM conventionally-scored REM and NREM sleep, is a quantifiable measure that reflects this process. Our findings are seemingly at odds with the functional significance currently assigned to standard sleep measures. For examples, several mathematical models have been developed based on the two process model of sleep regulation^52^, proposed by Borbely (1982), consisting of Process S, a homeostatic pressure to enter NREM slow-wave activity (SWA) (which is thought to be the most recuperative type of sleep) and Process C, representing the circadian rhythm of alertness. At any given timepoint, the models assume that sleep propensity (sleepiness) is mainly a function of the combined influence of these two processes. Process S is generally assumed to uniformly increase with wake duration and decay (reduce sleep debt) with sleep. In other words, all sleep is assumed to be recuperative. A subclass of these models, mathematical performance prediction models (MPPMs), have been developed to predict daytime alertness/cognitive performance.

Many of the MPPMs predict some aspects of human performance reasonably well^35,36,53^; however, tests of the models are mostly conducted under laboratory conditions where there are severe restrictions on sleep opportunities that presumably limit the quantity of SWA, and thus the ability to recover from sleep debt accrued during the prior wakefulness period. With the data utilized for the present study, by design, such prior sleep debt was mostly accounted for as evidenced by the lack of sleep duration or SWA predictive value despite large variability in these measures. Our work does not rule out an important role for sleep duration and SWA as factors for predicting performance when sleep opportunity is limited but highlights the importance of REM-like activity in predicting performance under more natural conditions of unrestricted sleep opportunity. Our results highlight that sleep is complex with periods of REM-like activity that appear to incur additional sleep debt. Accordingly, it is likely that accounting for the effects of REM OTA in MPPMs and other sleep-wakefulness outcome models would enhance their predictive power and utility.

As noted, in sharp contrast to standard sleep measures, REM-like OTA showed a substantial positive correlation with individual differences in subsequent waking impaired-performance. This was likely aided (made salient) by the fact that the EEG sleep data was collected from subjects who were essentially sleep satiated – thus minimizing the extent to which interindividual differences in homeostatic sleep pressure could mask the influence of REM OTA. Currently unknown is the extent to which individual differences in sensitivity/resilience to sleep loss persist across a range of reduced sleep opportunities^54^. In other words, what is the basis of individual differences (IDs) in neurobehavioral sleep debt that persist after accounting for imposed sleep loss? These performance IDs may be due to factors unrelated to sleep, per se. The present findings, though, suggest that these differences may, in large part, be explained by individual differences in REM-like sleep. Within this vein the present findings are relevant to studies of individual differences in *resilience* under various sleep loss scenarios. Chua et al. (2019) argue that, prior to their work, there was no simple way to classify the resilience of individuals without subjecting them to severe sleep loss. They then showed that baseline (rested) differences in performance on the PVT can be used to classify individuals into one of three resilience bins^46^. Our findings indicate that REM-like OTA substantially predicts differences in baseline PVT effects, and as such is relevant to understanding how different individuals respond to sleep loss or how sleep loss may impact the same individual across days, weeks, months, etc.

REM-like OTA during sleep as found here accounted for 25-38% of the ID in PVT performance. This finding was consistent across data sets from two different studies. Remarkably, these studies had important differences in their design resulting in both different sleep and PVT statistics (see Table S1). Yet the REM-like OTA association with vigilance was clearly evident in both datasets. In addition, subsequent to the initial presentation of the present findings (in poster form)^55^, Ujma et al. (2023)^56^ reported related findings from a broad data-mining study. Despite the absence of a specific REM hypothesis, they reported significant albeit weak correlations between scored-REM oscillatory theta and performance on the Modified Mini Mental State Test, Trails B, and Digit Vigilance. Their analyses revealed negative correlations ranging from -0.05 to -0.1 (reduced performance). They also referenced a previous data-mining study in which similar results were obtained using an IQ measure^57^. Although these correlations were relatively weak (possibly reflecting, in part, the likelihood that the performance measures they utilized were less sensitive to variations in sleep debt than the PVT^34^), their findings are generally consistent with ours.

The present findings have several implications. If our hypothesis is correct, a seemingly paradoxical consequence would be that REM sleep may counteract the recuperative effects of the NREM sleep that dominates during the first half of the nocturnal sleep period. Why would this be the case? As hypothesized by others,^7,58,59^ the important putative functions of REM sleep (e.g. emotional regulation, synaptic plasticity, memory consolidation) for wakefulness may require, or benefit from, some sort of “priming” by NREM sleep and these REM sleep functions may be resource-intensive (i.e., utilize a portion of the restorative resources that were gained during the preceding period of NREM sleep such as metabolite clearance). The magnitude of performance deficits among healthy individuals with higher REM-like OTA power observed in the present study may not be severe enough to exert a meaningful effect on cognitive functions during typical daily life and are presumably significantly outweighed by the beneficial effects. However, the impact of high levels of REM-like OTA on information processing may reach pathological levels for certain predisposed individuals (e.g. neurological and psychiatric illnesses) and under specific conditions, such as high levels of stress or in those with cognitively demanding occupations. REM-like cognitive neurophysiological activity should be investigated in these populations using similar OTA-type measures throughout sleep. Importantly, as suggested by our results, this can and should be done without strict regard to conventional sleep stage scoring criteria as REM-like neural activity may be transformed (deviate from traditional scoring but still be elevated) vs reduced in these populations.

The present results suggest the potential utility of reanalyzing EEG data from prior studies using a more neurophysiology-specific measure of REM-like sleep processes. For example, our comparison of different measures of REM activity (see Figure 2), suggests that previous REM EEG/theta analyses based on conventionally scored REM sleep likely underestimate the effect size of REM neural activity and were more susceptible to noise due to poorer signal-to-noise ratios. Utilizing the oscillatory theta power (with background 1/f theta power and other sources of OTA removed) and employing a less (conventional, manual-scoring) biased nOTA measure throughout sleep should yield stronger and more reliable predictions of subsequent wakefulness effects. For example, in a hypothetical study comparing REM theta power in a patient group versus a control group, a very high 1/f theta power may render a meaningful difference in oscillatory theta power difficult or impossible to detect if it is not removed from the analysis. In addition to REM-OTA being a stronger predictor, its nOTA counterpart offers the advantage of being amenable to automation, eliminating the need for human scorers or machine training on conventionally-scored data. nOTA is a superior measure compared to scored-REM raw theta activity, the current standard approach, since it captures both the duration and amplitude of the entire night of REM-like OTA without the relatively arbitrary constraints that are imposed by conventional sleep scoring.

It is possible that our finding is strictly a trait-like association that lacks a (near-time direct) causal mechanism, and is due to some third factor that interferes with performance. Although our findings are consistent with the *REM-OTA Fatigue Hypothesis*, future work should be focused on longitudinal data across several days/nights to assess intra-individual variation, as a further test of the present hypothesis. REM manipulation studies are also warranted, though these should use alternative REM-like measures to properly account for changes in REM-like neural activity that may only loosely vary as a function of conventionally scored sleep stages. In other words, effort should be made to discriminate a change in how REM-like neural activity presents vs reduction/increase with a REM intervention when testing this hypothesis further.

Acknowledging limitations of interpreting the work presented here, given what we know about the biology of REM-like OTA, we strongly suspect that our findings do in fact reflect causal mechanisms. In addition to the present results, perusal of the relevant scientific literature reveals a number of incidental findings that tend to confirm the REM OTA hypothesis. A transgenic mouse model study in which the primary objective was to identify and characterize REM regulatory cells, revealed that subsequent to natural and induced increases in REM duration there was a concomitant increase in NREM slow-wave activity (0.5-4 Hz, SWA)^60^, presumably the recuperative process for sleep debt. In addition, a REM sleep deprivation experiment in humans revealed longer sleep latencies with multiple sleep latency tests^61^ potentially reflecting less sleep pressure (debt) and another found decreased SWA band power^62^. These findings suggest that sleep debt is reduced or increased when REM sleep is artificially reduced or increased, respectively. While these intervention studies tend to support the hypothesis that the drive for NREM recuperative processes accrues as a function of REM processes, these findings may be artifactual effects of the interventions. However, numerous natural sleep studies (although lacking post-sleep measures of wakefulness and performance) in which relative NREM and REM durations were assessed are also consistent with the REM OTA hypothesis. They show positive correlations between scored- NREM durations and previous scored-REM durations across species, and rarely the converse (rodent^60,63,64^, cat^65^, non-human primate^66^, and humans^67^). Thus, there is considerable evidence that REM-like neural activity incurs additional sleep debt, and as we have found, that this additional sleep debt is evident and measurable during subsequent wakefulness. While on whole sleep may be recuperative, net recovery appears to be a function of the balance between sleep processes that restore and those that incur a cost. This suggests that a significant reevaluation of current assumptions regarding recuperation during sleep is warranted, as well as follow-up investigations focused on the REM OTA hypothesis and its implications.

## Author Contributions

SV and JDH conceived of and implemented the analyses. AB and TB contributed critical feedback for the analytical approach. SV, JDH, and TB wrote the manuscript. TJD, SRM, and AB contributed to editing the manuscript.

## Acknowledgements

We thank Carson C Chow for his input on the psychomotor vigilance task generative model.

## Disclaimer

Material has been reviewed by the Walter Reed Army Institute of Research. There is no objection to its presentation and/or publication. The opinions or assertions contained herein are the private views of the authors, and are not to be construed as official, or as reflecting true views of the Department of the Army or Department of Defense. The investigators have adhered to the policies for protection of human subjects as prescribed in AR 70-25.

## Data

We combined data from two human sleep research laboratory studies. Participants were 47 healthy young adults screened for sleep disorders and health problems (Study A n = 26, Study B n = 21). Two exclusion criteria based on sleep staging and spectral quality were used: 1) long sleep onset latency (SOL) of 75 minutes for Study A and 38 minutes for Study B (defined as the first occurrence of 3 epochs of sleep), and 2) high variance of the time-frequency spectrogram across the low-bands (0.5-15Hz) due to residual noise after Luna QC (defined below). Exclusion thresholds were determined using standard group-wise statistical cutoffs from Tukey’s method. An outlier was anyone greater than 1.5 x IQR. Five out of 47 original subjects were removed based on these criteria, resulting in 24 subjects from Study A (16 female, median age 24 [range: 18 to 33 years]), and 18 subjects from Study B (7 female, median age 24 [range: 19 to 33 years]).

To help ensure that participants entered the laboratory with minimal sleep debt, participants were asked to maintain an 8-hour sleep schedule while at-home for at least 7 days. In Study A this was followed by three baseline nights in the sleep laboratory with 10-hour TIB each night. In Study B, the baseline was one 8-hour nighttime in-laboratory sleep period. Only EEG data from the last or only baseline in- laboratory sleep opportunity were included in the analyses. As outlined below, on the day following the last baseline night, performance testing was conducted at regular intervals (Study A, 75 minutes or Study B, 120-minute intervals) using the Psychomotor Vigilance Test (PVT). An average PVT reaction time using a robust estimator (see below for details) was inferred for the performance of each subject.

## Methods

### Polysomnography (PSG) main processing and predictors

Polysomnographic data was collected in both studies. A single Fz-channel with mastoid reference, sampled at 256 Hz, was used in both. PSG was acquired with the Natus Neuroworks system. The Luna sleep analysis platform^1^ was used for EEG artifact detection across 30-second epochs. The artifact detection algorithm consisted of: 1) bandpass filter from 0.1 to 128 Hz (Kaiser window method: ripple = 0.02, transition width = 1), 2) artifact detection using the Buckelmueller et al. method^1^, and 3) Hjorth parameters^1^ using iterative thresholding (three iterations) based on 3 standard deviations from mean Hjorth estimates per-epoch. These filters were either used to generate new EDF files for each conventionally scored stage or to generate epoch-mask files to be used for more fine-grained analyses.

Our approach requires the isolation of REM-like oscillatory theta-band activity from both aperiodic broad-band activity and other sources of oscillatory theta that obscure this signal. Theta-band activity was defined as as oscillatory signals in the 4-8 Hz band. (We did note variation across individuals in their peak frequencies but found that this fixed range was sufficient for our analyses.) Aperiodic EEG activity refers to the well-known broad-band activity that dominates EEG recordings and that approximately scales with the inverse of spectral frequency due to multiple sources of activity and filtering^2^. As shown in Figure 5, this activity dominates the theta-band, necessitating the utilization of additional methods to isolate the REM-like oscillatory theta activity.

Recently, several strategies for isolating oscillatory-type activity from 1/f-type spectra have been investigated. Two popular methods are: (a) a curve-fitting generalized linear model (e.g., fitting oscillations and one over f, FOOOF) or (b) resampling methods. Resampling was chosen for our analyses. We found similar early results using FOOOF. We used a resampling method called irregular resampling auto-spectral analysis (IRASA). Resampling attempts to isolate the aperiodic activity by leveraging a fractal hypothesis. For a fractal time series (self-affine property)^3^, up/down resampling (how often the time series is sampled) of the time series by *h* results in a similar time series, where the resulting new spectra are proportional with scaling by the rate *h* and Hurst exponent (*H*),

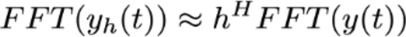

In principle, while the fractal component will remain intact, the oscillatory (periodic) components of the series will be shifted due to the resampling. Roughly, the IRASA method uses reciprocal pairs of sampling rates to displace an oscillatory peak down and up in frequency band and then takes the element- wise product of these two spectra (theoretically attenuating the shifted oscillatory peaks) resulting in a *cross-spectra*. This is repeated for several pairs of *(h*, *h^-^*^1^), taking the median of these cross-spectra, further attenuating oscillatory peaks and resulting in an inferred *aperiodic spectrum*. The *oscillatory spectrum* is then estimated by subtracting the aperiodic from the raw (original) power spectrum.

Specifically in the presented work, for each time series (concatenated time series across the whole sleep period for a specific scored-stage or 30-second epochs independent of scoring), we generated 5 IRASA pairs using upsample factors - h ∈ {1.01, 1.13, 1.26 , 1.38, 1.50} - and their reciprocals h^-1^. Welch’s method using a median estimator and hamming window was then applied to the original and resampled time series. We used a 10-second window duration for Welch’s method resulting in 0.1 Hz spectral bins. For better visualization, 1-second windows resulting in 1 Hz spectral bins were used for Figure 6

We incorporated the IRASA method into a time-frequency (spectrogram) algorithm, estimating oscillatory power across 30-second epochs. This required additional processing to reduce spurious noise effects due to finite sampling. For this, we used median and Gaussian smoothing. Median smoothing was 0.5 Hz across bands and 7.5 minutes across time. Gaussian standard deviation was 0.8 Hz and 4 minutes. (Note that theoretically, we could use shorter epochs than 30-second but this would introduce more noise due to finite sampling.)

For a given power band of interest, a predictor consisted of the sum over Welch-derived median bins in this range (and the equivalent average over sums for the IRASA spectrogram derived predictors). Power bands consisted of: theta (4-8Hz) and slow-wave activity (0.5-4Hz).

### Normalization of IRASA-derived spectrograms

For AASM agnostic spectrogram analyses, we found that using a normalization factor based on activity in the 0.5-35 Hz range improved prediction. This was based on the hypothesis that REM-like oscillatory theta-band activity when it occurs will be the dominant activity for frequencies less than 35 Hz. Reasons for this range were specifically to reduce slow-wave activity (SWA) sources of OTA (e.g. slow oscillation (SO) associated theta-bursts that are presumably unrelated to REM-like OTA) and wakefulness alpha- band activity (8-12Hz) (which can occur during sleep as well). The normalization was done at each 30- second epoch since the purpose was to isolate the source of OTA at each epoch. The normalization was implemented by dividing each time-frequency element of the smoothed IRASA-derived spectrogram (details above) by the normalization factor for a respective epoch. To quantify this normalization factor, we used the maximum power within the 0.5-35Hz band. This rescales the oscillatory spectra such that the dominant process will be peaked at 1. For example, even if there is a SO-associated OTA it will be dwarfed by the SO peak (relatively miniscule SO-OTA compared to SO power).

### Conjugate eye-movement measure

During wakefulness and REM sleep, there is coordination between the eyes to fixate on a single object (aka *conjugate eye movement*). We used this to visualize how well our REM-like neurophysiology measures separated wakefulness from REM-like sleep (since conjugate eye movement would be thought to be prominent but non-specific to wakefulness and REM sleep). We developed a conjugate eye movement measure to help track both wake and REM sleep (or conversely NREM sleep) based on the fact that left and right lateral rectus muscles (extraocular muscle channels) are anti-correlated during these periods. The measure was calculated as:

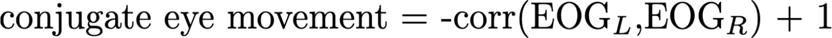

### Psychomotor vigilance task (PVT) task design

In Study A a visual computer-based 10-minute PVT^4^ was administered every 75 minutes during post baseline night waking period (interstimulus interval [ISI]: 2-10 seconds, with a uniform distribution). In Study B, a 5-minute PVT was administered every two hours (ISI: 1-5 seconds, with a uniform distribution). The 10-minute version was associated with larger group-wise mean reaction time (RT) and variance (mean=319 msec) than the 5-minute version (mean=278 msec). To address possible confounding by the different study-specific PVT statistics, data were z-scored prior to combining the two independent data sets.

### Vigilance data and robust mean reaction time estimates

For each subject, we estimated their mean PVT reaction time (PVT-RT) by aggregating data across sessions and using the following robust statistic to reduce spurious effects and for more advanced modeling. Mean reaction times (RT) were calculated from a generative (biophysical) Bayesian model. PVT response times were hypothesized to be drawn from a mixture of processes: the physiological processing of cue-and-motor response and attentional lapses. As such, in the model, we assumed that the data were derived from a mixture of a gamma distribution for response times that is consistent with a multistage process, and attentional lapses modeled as a uniform distribution, given by:

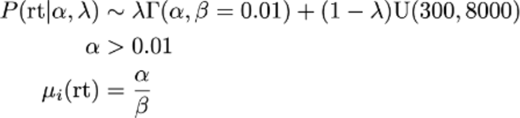

Parameters were inferred using the Stan platform and Hamiltonian Monte Carlo method (No-U-Turn sampler)^2^. The mean PVT-RT is given by the variable μ. False starts defined as both premature responses (i.e., responding when there is not a stimulus) and coincidental responses (i.e., response times less than 100 ms) were removed before calculating means. Our results held when using a simpler arithmetic mean but were noisier in some cases. The Bayesian method was also preferred for further generative modeling in our other work.

PVT-sleep measure correlations were estimated using the Python statsmodel package^5^. Data from each study (exogenous and endogenous variables) were z-scored and then aggregated. Ordinary least squares regression was then performed on these aggregated data. P-values were estimated using the Python statsmodel method, and two-tailed p-values are reported for t-scores.

### Code

The IRASA spectrogram code with examples is available at: https://github.com/ShashaankV/irasa_spectrogram

### Data Availability

Data is available through request and appropriate Data Use Agreement.

## Supplementary Materials

**Table S1.** Duration estimates (TST, SOL, WASO, NREM, REM) based on AASM conventionally scored 30-second epochs. Total sleep time estimated using the number of 30-second epochs scored as NREM 1, 2, or 3 based on conventional AASM criteria. SOL based on the first occurrence of three consecutive sleep epochs.

| Measure | Study A<br>mean (range) | Study B<br>mean (range) |
| --- | --- | --- |
| **Demographics** |  |  |
| Subjects | 24 | 18 |
| Female | 16 | 7 |
| Age (years) median (range) | 24 (18-33) | 24 (19-33) |
| **Sleep Statistics** |  |  |
| Prior in-lab satiation days | 2 | 0 |
| Time in Bed (TIB) (hours) | 10 | 8 |
| Total Sleep Time (TST) (hours) | 8.6 (6.4-9.6) | 7 (5.6-7.8) |
| Sleep Onset Latency (SOL)<br>(minutes) | 22.1 (4-61) | 11.3 (3-29.5) |
| Wake After Sleep Onset<br>(WASO) | 67.9 (14.5-192.5) | 45.2 (8-112.5) |
| NREM Stage 3 (% TST) | 17 (9-29) | 18 (4-32) |
| NREM Stage 2 (% TST) | 52 (39-60) | 53 (33-78) |
| REM (% TST) | 22 (14-33) | 19 (12-29) |
| **PVT Statistics** |  |  |
| PVT ISI (seconds) | 5 (2-10) | 2.5 (1-5) |
| PVT session duration (minutes) | 10 | 5 |
| Inter-individual differences in<br>mean PVT-RT (milliseconds) | 319 (273-392) | 278 (250-311) |

## References

1. Vyazovskiy, V. V. Sleep, recovery, and metaregulation: explaining the benefits of sleep. Nat. Sci. Sleep 7, 171–184 (2015).

2. Krueger, J. M., Frank, M. G., Wisor, J. P. & Roy, S. Sleep function: Toward elucidating an enigma. Sleep Med. Rev. 28, 46–54 (2016).

3. Obal, F. & Krueger, J. M. Biochemical regulation of non-rapid-eye-movement sleep. Front. Biosci. J. Virtual Libr. 8, d520–550 (2003).

4. Mergenthaler, P., Lindauer, U., Dienel, G. A. & Meisel, A. Sugar for the brain: the role of glucose in physiological and pathological brain function. Trends Neurosci. 36, 587–597 (2013).

5. Jones, B. E. Glia, Adenosine, and Sleep. Neuron 61, 156–157 (2009).

6. Renouard, L. et al. REM sleep promotes bidirectional plasticity in developing visual cortex in vivo. Neurobiol. Sleep Circadian Rhythms 12, 100076 (2022).

7. Zhou, Y. et al. REM sleep promotes experience-dependent dendritic spine elimination in the mouse cortex. Nat. Commun. 11, 4819 (2020).

8. Miller, K. E. & Gehrman, P. R. REM Sleep: What Is It Good For? Curr. Biol. 29, R806–R807 (2019).

9. Bandarabadi, M. et al. Dynamic modulation of theta–gamma coupling during rapid eye movement sleep. Sleep 42, (2019).

10. Vijayan, S., Lepage, K. Q., Kopell, N. J. & Cash, S. S. Frontal beta-theta network during REM sleep. eLife 6, e18894 (2017).

11. Armitage, R. The distribution of EEG frequencies in REM and NREM sleep stages in healthy young adults. Sleep 18, 334–341 (1995).

12. Bergel, A., Deffieux, T., Demené, C., Tanter, M. & Cohen, I. Local hippocampal fast gamma rhythms precede brain-wide hyperemic patterns during spontaneous rodent REM sleep. Nat. Commun. 9, 5364 (2018).

13. Mishra, A. & Colgin, L. L. The High Energy Cost of Theta–Gamma Activity during REM Sleep. Trends Neurosci. 42, 239–241 (2019).

14. Pal, D. & Mashour, G. A. Consciousness, Anesthesia, and Acetylcholine. Anesthesiology 134, 515–517 (2021).

15. Newman, E. L., Gupta, K., Climer, J. R., Monaghan, C. K. & Hasselmo, M. E. Cholinergic modulation of cognitive processing: insights drawn from computational models. Front. Behav. Neurosci. 6, 24 (2012).

16. Hutchison, I. C. & Rathore, S. The role of REM sleep theta activity in emotional memory. Front. Psychol. 0, (2015).

17. Lee, M. G., Hassani, O. K., Alonso, A. & Jones, B. E. Cholinergic Basal Forebrain Neurons Burst with Theta during Waking and Paradoxical Sleep. J. Neurosci. 25, 4365–4369 (2005).

18. Lechat, B. et al. New and Emerging Approaches to Better Define Sleep Disruption and Its Consequences. Front. Neurosci. 15, (2021).

19. Nielsen, T. A. A review of mentation in REM and NREM sleep: ‘covert’ REM sleep as a possible reconciliation of two opposing models. Behav. Brain Sci. 23, 851–866; discussion 904-1121 (2000).

20. Bernardi, G. et al. Regional Delta Waves In Human Rapid Eye Movement Sleep. J. Neurosci. 39, 2686–2697 (2019).

21. Nazari, M. et al. Regional variation in cholinergic terminal activity determines the non-uniform occurrence of cortical slow waves during REM sleep in mice. Cell Rep. 42, (2023).

22. Wang, Z. et al. REM sleep is associated with distinct global cortical dynamics and controlled by occipital cortex. Nat. Commun. 13, 6896 (2022).

23. Vattikuti, S. et al. Oscillatory Theta-Band Activity as a Sleep Stage Independent Measure of REM-like Activity throughout Sleep. Sleep 45, A42–A43 (2022).

24. Cox, R., Rüber, T., Staresina, B. P. & Fell, J. Phase-based coordination of hippocampal and neocortical oscillations during human sleep. *Commun*. Biol. 3, 176 (2020).

25. Boyce, R., Glasgow, S. D., Williams, S. & Adamantidis, A. Causal evidence for the role of REM sleep theta rhythm in contextual memory consolidation. Science 352, 812–816 (2016).

26. Kocsis, B. REMembering what you learned. Science 352, 770–771 (2016).

27. Borst, J. G. G., Leung, L.-W. S. & MacFabe, D. F. Electrical activity of the cingulate cortex. II. Cholinergic modulation. Brain Res. 407, 81–93 (1987).

28. Cantero, J. L. et al. Sleep-Dependent θ Oscillations in the Human Hippocampus and Neocortex. J. Neurosci. 23, 10897–10903 (2003).

29. He, B. J., Zempel, J. M., Snyder, A. Z. & Raichle, M. E. The Temporal Structures and Functional Significance of Scale-free Brain Activity. Neuron 66, 353–369 (2010).

30. Belluscio, M. A., Mizuseki, K., Schmidt, R., Kempter, R. & Buzsáki, G. Cross-frequency phase-phase coupling between θ and γ oscillations in the hippocampus. J. Neurosci. Off. J. Soc. Neurosci. 32, 423–435 (2012).

31. Tononi, G. & Cirelli, C. Sleep and the Price of Plasticity: From Synaptic and Cellular Homeostasis to Memory Consolidation and Integration. Neuron 81, 12–34 (2014).

32. Gottesmann, C. The transition from slow-wave sleep to paradoxical sleep: evolving facts and concepts of the neurophysiological processes underlying the intermediate stage of sleep. Neurosci. Biobehav. Rev. 20, 367–387 (1996).

33. Wen, H. & Liu, Z. Separating Fractal and Oscillatory Components in the Power Spectrum of Neurophysiological Signal. Brain Topogr. 29, 13–26 (2016).

34. Balkin, T. J. et al. Comparative utility of instruments for monitoring sleepiness-related performance decrements in the operational environment. J. Sleep Res. 13, 219–227 (2004).

35. Rajdev, P. et al. A unified mathematical model to quantify performance impairment for both chronic sleep restriction and total sleep deprivation. J. Theor. Biol. 331, 66–77 (2013).

36. Dongen, H. P. A. V., Dongen, V. & Comparison, H. Comparison of mathematical model predictions to experimental data of fatigue and performance. Aviat. Space Environ. Med. 15–36 (2004).

37. Van Dongen, H. P. A., Maislin, G., Mullington, J. M. & Dinges, D. F. The Cumulative Cost of Additional Wakefulness: Dose-Response Effects on Neurobehavioral Functions and Sleep Physiology From Chronic Sleep Restriction and Total Sleep Deprivation. Sleep 26, 117–126 (2003).

38. Honn, K. A. et al. New insights into the cognitive effects of sleep deprivation by decomposition of a cognitive throughput task. Sleep 43, zsz319 (2020).

39. Belenky, G. et al. Patterns of performance degradation and restoration during sleep restriction and subsequent recovery: a sleep dose-response study. J. Sleep Res. 12, 1–12 (2003).

40. Mollicone, D. J., Van Dongen, H. P. A., Rogers, N. L., Banks, S. & Dinges, D. F. Time of day effects on neurobehavioral performance during chronic sleep restriction. Aviat. Space Environ. Med. 81, 735–744 (2010).

41. McCauley, M. E. et al. Fatigue risk management based on self-reported fatigue: Expanding a biomathematical model of fatigue-related performance deficits to also predict subjective sleepiness. Transp. Res. Part F Traffic Psychol. Behav. 79, 94–106 (2021).

42. Reifman, J., et al. 2B-Alert App: A mobile application for real-time individualized prediction of alertness. J. Sleep Res. 28, e12725 (2019).

43. Arsintescu, L., Kato, K. H., Hilditch, C. J., Gregory, K. B. & Flynn-Evans, E. Collecting Sleep, Circadian, Fatigue, and Performance Data in Complex Operational Environments. JoVE J. Vis. Exp. e59851 (2019) doi:10.3791/59851.

44. Huang, Y., Hennig, S., Fietze, I., Penzel, T. & Veauthier, C. The Psychomotor Vigilance Test Compared to a Divided Attention Steering Simulation in Patients with Moderate or Severe Obstructive Sleep Apnea. Nat. Sci. Sleep 12, 509–524 (2020).

45. Zhang, C. et al. Psychomotor Vigilance Testing of Professional Drivers in the Occupational Health Clinic: a Potential Objective Screen for Daytime Sleepiness. J. Occup. Environ. Med. Am. Coll. Occup. Environ. Med. 54, 296–302 (2012).

46. Chua, E. C.-P. et al. Classifying attentional vulnerability to total sleep deprivation using baseline features of Psychomotor Vigilance Test performance. Sci. Rep. 9, 12102 (2019).

47. Chua, E. C.-P. et al. Sustained Attention Performance during Sleep Deprivation Associates with Instability in Behavior and Physiologic Measures at Baseline. Sleep 37, 27–39 (2014).

48. Huber, R. et al. Human Cortical Excitability Increases with Time Awake. Cereb. Cortex 23, 1–7 (2013).

49. Finelli, L. A., Baumann, H., Borbély, A. A. & Achermann, P. Dual electroencephalogram markers of human sleep homeostasis: correlation between theta activity in waking and slow-wave activity in sleep. Neuroscience 101, 523–529 (2000).

50. Hughes, J., Doty, T. J., Ratcliffe, R. & Balkin, T. J. Slow oscillatory transcranial direct current stimulation during sleep enhances cognitive performance during subsequent sleep deprivation. Sleep 44, (2021).

51. Gonzalez, C. E. et al. Theta Bursts Precede, and Spindles Follow, Cortical and Thalamic Downstates in Human NREM Sleep. J. Neurosci. Off. J. Soc. Neurosci. 38, 9989–10001 (2018).

52. Achermann, P. & Borbély, A. A. Mathematical models of sleep regulation. Front. Biosci.- Landmark 8, 683–693 (2003).

53. Johnson, M. et al. Modulating the Homeostatic Process to Predict Performance during Chronic Sleep Restriction. Aviat. Space Environ. Med. 75, A141–6 (2004).

54. Rupp, T. L., Wesensten, N. J. & Balkin, T. J. Trait-Like Vulnerability to Total and Partial Sleep Loss. Sleep 35, 1163–1172 (2012).

55. Vattikuti, S. et al. REM-like Neural Activity Is Superior to NREM Parameters for Predicting Non-sleep Restricted Vigilance. Sleep 45, A43–A44 (2022).

56. Ujma, P. P. et al. Multivariate prediction of cognitive performance from the sleep electroencephalogram. 2023.02.28.530401 Preprint at 10.1101/2023.02.28.530401 (2023).

57. Ujma, P. P. et al. The sleep EEG spectrum is a sexually dimorphic marker of general intelligence. Sci. Rep. 7, 18070 (2017).

58. Benington, J. H. & Heller, H. C. Does the function of REM sleep concern non-REM sleep or waking? Prog. Neurobiol. 44, 433–449 (1994).

59. Llewellyn, S. & Hobson, J. A. Not only…but also: REM sleep creates and NREM Stage 2 instantiates landmark junctions in cortical memory networks. Neurobiol. Learn. Mem. 122, 69–87 (2015).

60. Hayashi, Y. et al. Cells of a common developmental origin regulate REM/non-REM sleep and wakefulness in mice. Science 350, 957–961 (2015).

61. Nykamp, K. et al. The Effects of REM Sleep Deprivation on the Level of Sleepiness/Alertness. Sleep 21, 609–614 (1998).

62. Beersma, D. G., Dijk, D. J., Blok, C. G. & Everhardus, I. REM sleep deprivation during 5 hours leads to an immediate REM sleep rebound and to suppression of non-REM sleep intensity. Electroencephalogr. Clin. Neurophysiol. 76, 114–122 (1990).

63. Benington, J. H. & Heller, H. C. REM-sleep timing is controlled homeostatically by accumulation of REM-sleep propensity in non-REM sleep. Am. J. Physiol. 266, R1992–2000 (1994).

64. Vivaldi, E. A., Ocampo, A., Wyneken, U., Roncagliolo, M. & Zapata, A. M. Short-term homeostasis of active sleep and the architecture of sleep in the rat. J. Neurophysiol. (1994) doi:10.1152/jn.1994.72.4.1745.

65. Ursin, R. Sleep stage relations within the sleep cycles of the cat. Brain Res. 20, 91–97 (1970).

66. Weitzman, E. D., Kripke, D. F., Pollak, C. & Dominguez, J. Cyclic Activity In Sleep of Macaca Mulatta. Arch. Neurol. 12, 463–467 (1965).

67. Barbato, G. & Wehr, T. A. Homeostatic Regulation of REM Sleep in Humans During Extended Sleep. Sleep 21, 267–276 (1998).

## References

1. Overview - Luna. http://zzz.bwh.harvard.edu/luna/.

2. He, B. J., Zempel, J. M., Snyder, A. Z. & Raichle, M. E. The Temporal Structures and Functional Significance of Scale-free Brain Activity. Neuron 66, 353–369 (2010).

3. Wen, H. & Liu, Z. Separating Fractal and Oscillatory Components in the Power Spectrum of Neurophysiological Signal. Brain Topogr. 29, 13–26 (2016).

4. Khitrov, M. Y. et al. PC-PVT: a platform for psychomotor vigilance task testing, analysis, and prediction. Behav. Res. Methods 46, 140–147 (2014).

5. Seabold, S. & Perktold, J. Statsmodels: Econometric and Statistical Modeling with Python. in 92–96 (2010). doi:10.25080/Majora-92bf1922-011.

